# Sodium Benzoate Promotes Fat Accumulation and Aging via the SKN-1/Nrf2 Signaling Pathway: Evidence from the *Caenorhabditis elegans* Model

**DOI:** 10.1101/2024.09.16.613358

**Authors:** Jiah D. Lee, Jiwoo Lee, Jerry Vang, Xiaoping Pan

**Author notes:** Corresponding author: Xiaoping Pan, Ph.D.

## Abstract

Sodium benzoate (SB) is widely used in food products, cosmetics, and medical solutions due to its antimicrobial properties. While it is generally considered safe and has potential neuroprotective benefits, SB has also been linked to adverse effects, including hepatic oxidative stress and inflammation. However, the potential effects of SB on obesity and aging remain poorly understood. In this study, we investigated the effects of SB on fat accumulation and aging using the nematode *Caenorhabditis elegans* (*C. elegans*) as a model system. Wild-type worms were exposed to various SB concentrations (0%, 0.0004%, 0.0008%, 0.004%, and 0.1%) and 0.016% glucose as a positive control for 72 hours in liquid or on NGM agar plates. Fat accumulation was assessed through the Oil Red O staining, which revealed that SB induced more fat accumulation compared to vehicle control, even at low concentrations, including the dosage of 0.0004%. Lifespan analysis also demonstrated that SB significantly accelerated aging in wild-type worms in a dose-dependent manner. Further investigations found that SKN-1 (an Nrf2 homolog) is necessary for SB-induced fat accumulation and aging. Moreover, SB inhibited the nuclear localization of SKN-1 under oxidative stress conditions. These findings suggest that SB may induce fat accumulation and aging by inhibiting the oxidative stress-mediated SKN-1 signaling pathway.

## INTRODUCTION

Sodium benzoate (SB) (**Figure 1A**), a widely used preservative, is currently subjected to scrutiny due to recent studies on the health risks associated with its consumption. Recent research proposes a possible connection between SB and obesity (Ciardi et al., 2012). As obesity rates continue to rise, there is an increasing need to identify environmental obesogens and establish preventative measures to address this significant health concern. According to the Centers for Disease Control and Prevention National Center for Health Statistics (CDC NCHS), in the United States (US), 42.4% of adults (ages ≥ 20) were diagnosed as obese in 2017-2018 (Hales et al., 2020), and 18.5% of youth (ages 2-19) were estimated to be obese (Hales et al., 2017). According to the World Health Organization (WHO) (Available from: https://www.who.int/news-room/fact-sheets/detail/obesity-and-overweight), the obesity rate nearly tripled in 2020 compared to 1975. In 2016, approximately 1.9 billion (39%) adults were overweight, with 650 million (13%) considered to be obese. Individuals classified as either overweight or obese are at higher risks to develop diabetes, sleep apnea, stroke, hypertension, coronary heart disease, and various types of cancers (NHLBI Obesity Education Initiative Expert Panel on the Identification, Evaluation, and Treatment of Obesity in Adults (US). Clinical Guidelines on the Identification, Evaluation, and Treatment of Overweight and Obesity in Adults: The Evidence Report. Bethesda (MD): National Heart, Lung, and Blood Institute; 1998 Sep. Available from: https://www.ncbi.nlm.nih.gov/books/NBK2003/). Despite ongoing efforts to curb the rise in obesity rates, the availability and consumption of supposedly ‘safe’ food today exerts a significant impact on human health and nutrition. Given that processed foods comprise roughly 75% of the diet in Western societies, the utilization of preservatives is a familiar practice. While numerous preservatives have been extensively researched for their obesogenic effects (Simmons et al., 2014), still some others, such as SB, that have not been adequately investigated (Simmons et al., 2014).

**Figure 1.**
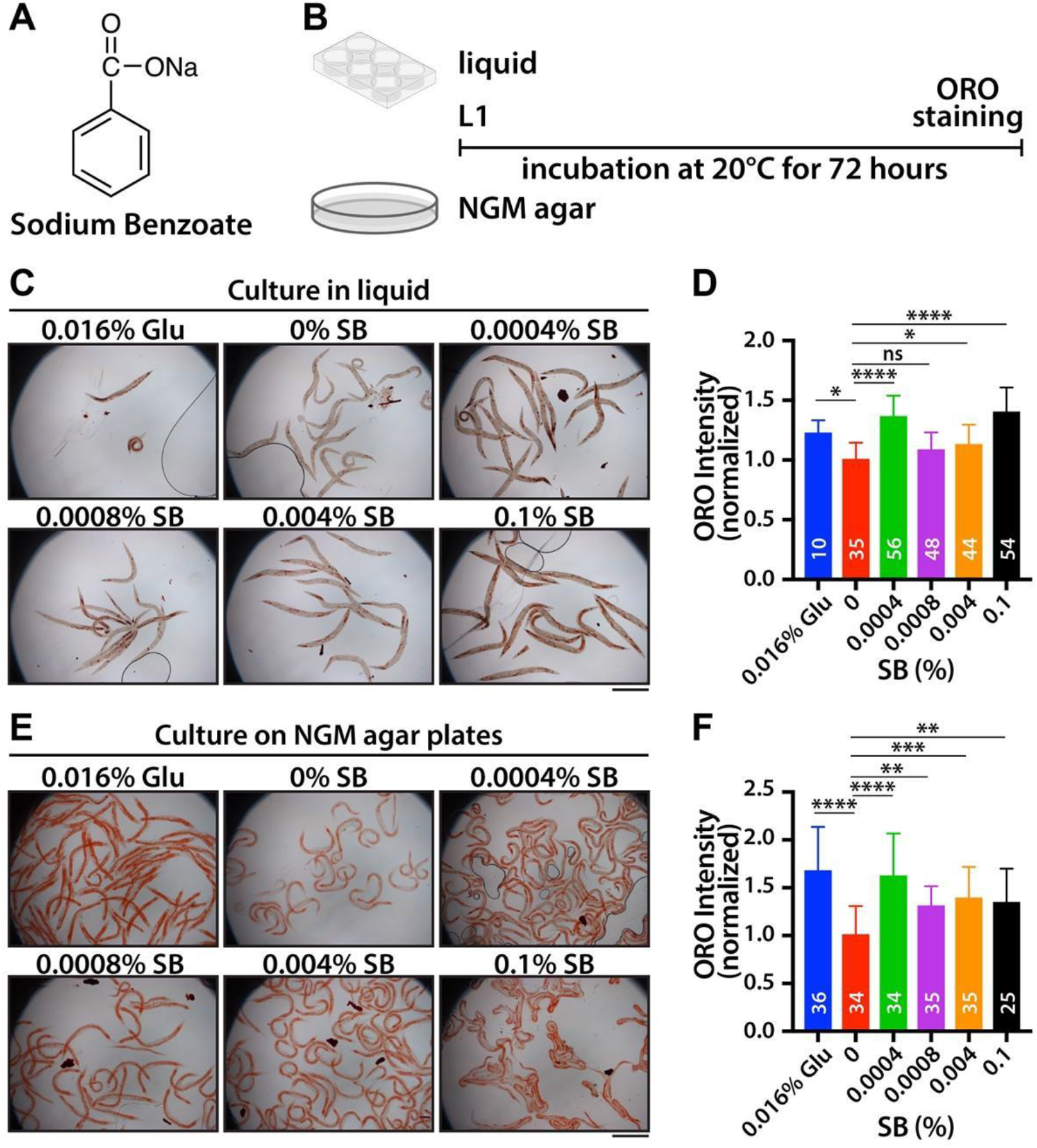
SB treatment increases fat accumulation in wild-type worms. (A) Chemical structure of sodium benzoate. (B) Strategy for SB treatment in liquid or on NGM agar plate. (C) ORO staining with SB-treated worms in liquid. (D) Normalized ORO intensity. (E) ORO staining with SB-treated worms on NGM agar plates. (F) Normalized ORO intensity. Not significant (ns), p > 0.05; *, 0.01 < p ≤ 0.05, **, 0.001 < p ≤ 0.01; ***, 0.0001 < p ≤ 0.001; **** p ≤ 0.0001. Scale bars: 500μm.

SB is currently classified as “Generally Regarded as Safe” (GRAS) by the US Food and Drug Administration (FDA), with a permitted concentration of up to 0.1% in food products (Lennerz et al., 2015). Recent studies have reported both positive and negative effects of SB on animal development and behaviors. For instance, in mice, SB has been found to improve cognitive functions following cortical impact injury (Rangasamy et al., 2021b) and stimulate dopamine production (Rangasamy et al., 2021a) but increase brain oxidative stress (Khoshnoud et al., 2018). Additionally, SB exposure delays development in fly larvae by altering endocrine hormone levels and commensal microbiota, and shorten their lifespan by inducing oxidative stress (Asejeje et al., 2023; Dong et al., 2022). SB also causes developmental defects, oxidative stress, and anxiety-like behaviors in zebrafish (Gaur et al., 2018). However, despite increasing studies on the toxicity of SB, few have explored its potential obesogenic effects (Ciardi et al., 2012). Therefore, investigating the obesogenic effects of SB could be crucial in addressing the issue of globally increasing obesity rates, particularly concerning the large-scale utilization of food preservatives.

The nematode *C. elegans* is a well-established model organism that has been used in a variety of studies due to its substantial genetic overlap with humans (60-80% homology) and conservation in biochemical and cellular pathways. It also features short generation and life cycles, self-fertilization, and transparency (Kaletta and Hengartner, 2006). Although *C. elegans* lacks leptin and designated fat tissues, it shares many aspects of similarities with higher organisms including humans, making it a valuable model organism for studying fat metabolism and obesity. For example, they store fat in the form of triglycerides in the intestines, oocytes, and hypodermis. They also share numerous conserved metabolic pathways regulating fat metabolism, such as the SKN-1 (an Nrf2 homolog) pathway (Pang et al., 2014). In this research, we investigated the effect of SB on obesity and aging, as these two are highly associated with each other (Shen et al., 2018). Wild-type worms were treated with various SB concentrations (0%, 0.0004%, 0.0008%, 0.004%, and 0.1%) and glucose (0.016%) as a positive control. Even at concentrations notably lower than the FDA’s permissible limit (0.1%), such as 0.0004%, SB induced significantly more fat accumulation and accelerated aging in wild-type worms as compared to control. Subsequent analysis using mutant worms revealed that SB-induced fat accumulation and aging are driven through the oxidative stress-mediated SKN-1 signaling pathway. Therefore, our findings suggest the potential risks of SB consumption with obesity and aging and a underlying mechanism and shed light on future studies in other organisms including humans.

## MATERIALS AND METHODS

### *C. elegans* culture and strains

*C. elegans* were maintained on nematode growth medium (NGM) agar plates seeded with *Escherichia coli* OP50 strain at 20°C (Brenner, 1974). The wild-type Bristol strain (N2) as well as mutant strains, including QV225 [*skn-1(zj15)*] (An and Blackwell, 2003) and LD1 [*idIs7 skn-1b/c::GFP + rol-6(su1006)*] (An and Blackwell, 2003), were provided by the *Caenorhabditis* Genetic Center (CGC).

### SB treatment

SB was purchased from Millipore Sigma (Cat#: PHR1231, MO, USA). To prepare a 5% stock solution, 2.5 g SB powder was dissolved in 50 mL distilled water. For the liquid SB treatment, the 5% SB stock solution was diluted with K-media (2.36 g KCl, 3 g NaCl, in 1 L ddH^*^O) to make final concentrations of 0%, 0.0004%, 0.0008%, 0.004%, and 0.1% SB in 24-well plates. For SB treatment on NGM agar plates, 5% SB stock solution was diluted into autoclaved NGM agar media (∼55°C) to make final concentrations of 0%, 0.0004%, 0.0008%, 0.004%, and 0.1% SB before being poured into 6 cm petri dish plates. L1-synchronized worms were cultured for 72 hours at 20°C in OP50-added K-media and on OP50-seeded NGM agar plates with various concentrations of SB and 0.016% glucose as a positive control. For starvation conditions, L4-synchronized worms were cultured for 72 hours at 20°C in K-media without OP50 *E. coli* but with various concentrations of SB and 0.016% glucose. For prolonged SB treatment, the first larval stage (L1)-synchronized worms were cultured for 120 hours at 20°C on OP50-seeded NGM agar plates with various concentrations of SB and 0.016% glucose.

### Oil Red O (ORO) staining

ORO staining was conducted as previously reported (Escorcia et al., 2018). The ORO stock solution was prepared by dissolving 500 mg of ORO powder (Cat#: O0625, Millipore Sigma, MO, USA) in 100 mL of 100% isopropanol. To prepare the ORO working solution, the ORO stock solution was diluted in water (3:2) to make a final concentration of 60% isopropanol. The ORO working solution was filtered using a 0.2 μm cellulose acetate sterile syringe filter. SB-treated worms were collected with 1×PBST (1×PBS plus 0.1% Tween 20) solution and washed three times with 1×PBST solution. After centrifugation, the supernatant was removed except for 100 μL, and 600 μL of 40% isopropanol was added to each worm pellet, followed by rocking at 20°C for 3 minutes. After centrifugation, the supernatant was removed except for 100 μL, and 600 μL of the ORO working solution was added to each sample. The samples were shaken in the dark for 2 hours at 20°C. All samples were washed six times for 1 hour with a 10-minute interval to remove excess ORO stain. Images of ORO-stained worms were captured using a Leica differential interference contrast (DIC) microscope equipped with a color camera.

### SKN-1::GFP expression analysis

To analyze the effect of SB on the localization of SKN-1::GFP in response to oxidative stress, L1-synchronized *skn-1b/c::gfp* transgenic worms were cultured for 48 hours at 20°C on OP50-seeded NGM agar plates with varying SB concentrations and 0.016% glucose. *skn-1b/c::gfp* transgenic worms were then collected using K-media and treated with 5 mM sodium arsenite for 30 minutes at 20°C to induce oxidative stress. To visualize the localization of SKN-1::GFP, *skn-1b/c::gfp* transgenic worms were dissected, fixed with 4% paraformaldehyde/1×PBS, and then post-fixed in cold methanol. The fixed intestines were incubated for 30 minutes with 1×PBST/0.5% BSA blocking solution, which consisted of 1×phosphate-buffered saline (PBS), 0.1% Tween 20, and 0.5% bovine serum albumin (BSA).

After washing three times, the dissected intestines were incubated for 2 hours at 25°C in antibodies against GFP (Abcam, Waltham, MA; Cat#: ab6556) and nuclear pore complex (NPC) proteins (mAb414, Abcam, Waltham, MA; Cat#: ab24609) diluted (1:500) in 1×PBST/0.5% BSA solution. After washing with 1×PBST/0.5% BSA solution three times, the dissected intestines were incubated for 2 hours at 25°C with secondary antibodies (ThermoFisher Scientific, Waltham, MA). The nuclei were stained with 4,6-diamidino-2-phenylindole (DAPI).

### Image analysis

The ORO intensities were measured using ImageJ software, with the intensity of each image subtracted from the background. The intensity of each sample was then normalized to the average intensity of 0% SB-treated worms per experiment. SKN-1::GFP intensities in the cytoplasm and nucleus were also measured using ImageJ software. The ratio of nuclear to cytoplasmic SKN-1::GFP proteins was calculated by dividing the nuclear SKN-1::GFP intensity by the cytoplasmic SKN-1::GFP intensity for each individual gonad.

### Lifespan analysis

L1-synchronized worms were cultured at 20°C on NGM agar plates with various concentrations of SB and 0.016% glucose, seeded with OP50 *E. coli*. All tested worms were transferred to fresh plates every three days, and their survival rates were scored daily.

### Statistical analysis

Statistical significance was analyzed using one-way analysis of variance (ANOVA) or the two-tailed Student’s t-test. The error bars represent the respective standard deviation (SD) values. The significance levels are denoted as follows: not significant (ns), p > 0.05; *, 0.01 < p ≤ 0.05; **, 0.001 < p ≤ 0.01; ***, 0.0001 < p ≤ 0.001; **** p ≤ 0.0001.

## RESULTS

### SB increases fat accumulation in wild-type worms

To evaluate whether SB exhibits an obesogenic effect, L1-synchronized wild-type worms were exposed to various concentrations of SB (0%, 0.0004%, 0.0008%, 0.004%, and 0.1%) and 0.016% glucose as a positive control for 72 hours in liquid (K-media) conditions. Fat accumulation was visualized by staining collected worms using Oil Red O (ORO) dye, as previously reported (Escorcia et al., 2018) (**Figure 1B**). The fat accumulation was primarily observed in the intestines, oocytes, and hypodermis. Fat accumulation levels in the entire body were quantified using ImageJ software and normalized by the ORO intensity of 0% SB-treated worms. A significant increase in fat accumulation was observed across the tested SB concentrations, except for 0.0008% of SB. Notably, higher fat accumulation was detected at 0.0004% and 0.1% SB (**Figures 1C and 1D**). To confirm this result, L1-synchronized wild-type worms were cultured on OP50-seeded NGM agar plates with varying SB concentrations and 0.016% glucose for 72 hours (**Figure 1B**). Fat accumulation levels were measured as described above. Similar patterns of fat accumulation were observed under NGM agar culture conditions (**Figures 1E and 1F**). Interestingly, 0.0004% SB treated worms had a higher fat accumulation than 0.0008% SB treated worms indicating a “low dose effect”. Our results suggest that SB increases fat accumulation in wild-type *C. elegans* worms even at much lower concentrations than the FDA’s permissible concentration (0.1%).

### SB may increase fat accumulation through food metabolism pathways

To explore the potential impact of SB on fat accumulation during starvation, fourth larval stage (L4)-synchronized wild-type worms were cultured in K-media with various concentrations of SB and 0.016% glucose in the absence of OP50 *E. coli* food. After 72 hours, the collected wild-type worms were stained with ORO dye. The results indicated no significant increase in fat accumulation (**Figure 2A**). Intriguingly, fat accumulation was even decreased at high SB concentrations (0.004% and 0.1%) (**Figure 2B**). Next, to investigate whether SB treatment enhances fat accumulation over an extended time, L1-synchronized wild-type worms were cultured for 120 hours on OP50-seeded NGM agar plates with varying SB concentrations and 0.016% glucose. It is known that aged worms typically exhibit an increasing deposit of fat. In this study we also observed that most aged adult worms (120 hours past L1) displayed more fat accumulation than young adult worms (72 hours past L1) (Compare **Figure 2C** to **Figure 1E**). Notably, prolonged SB treatment did not further increase fat accumulation compared to worms treated with 0% SB (**Figures 2C and 2D**). Therefore, these results indicate that SB treatment may enhance fat accumulation through food metabolic pathways which was also related to aging.

**Figure 2.**
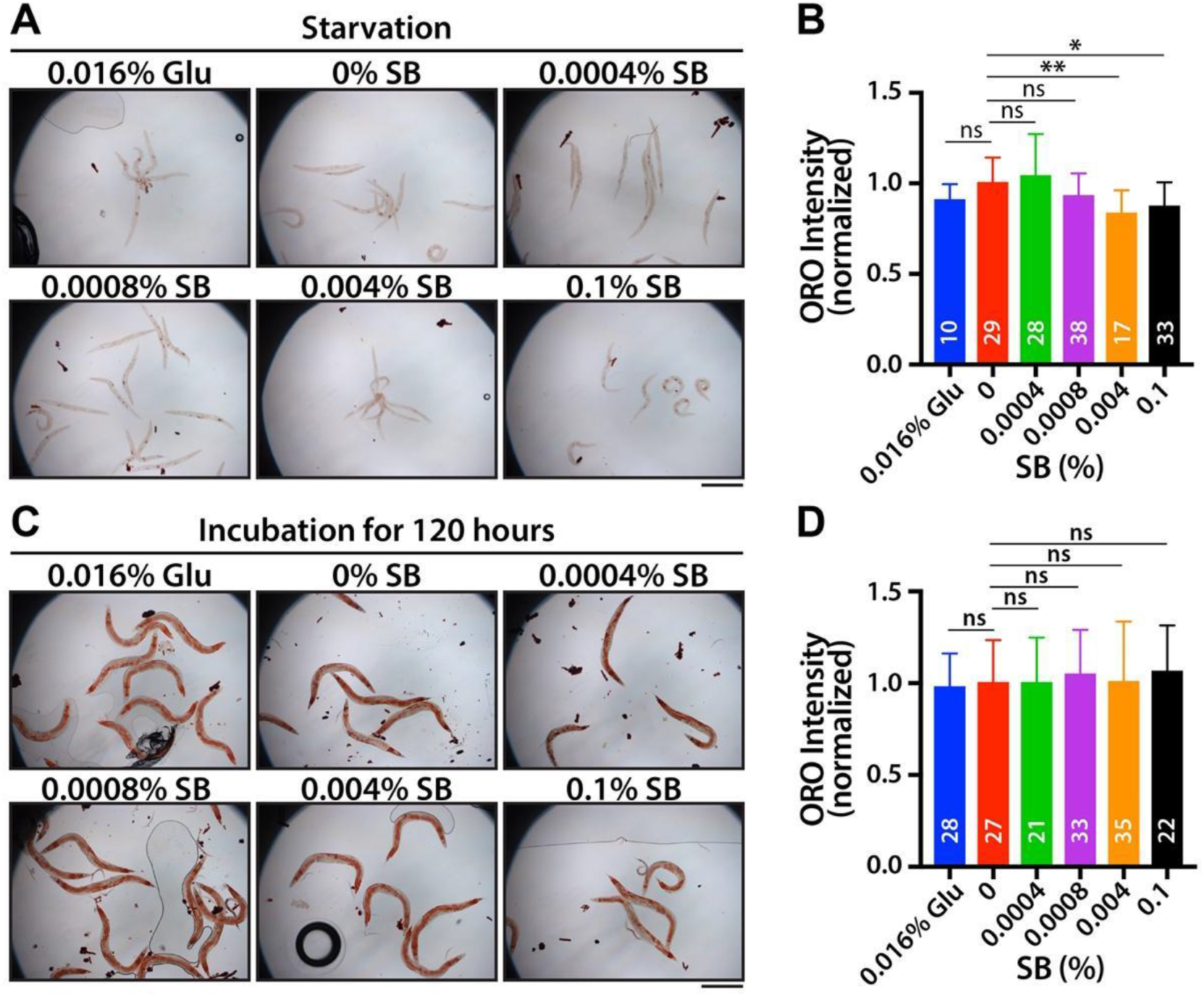
SB-induced fat accumulation may require metabolic pathways. (A) ORO staining with starved worms. (B) Normalized ORO intensity. (C) ORO staining with prolonged SB-treated worms. (D) Normalized ORO intensity. Not significant (ns), p > 0.05; *, 0.01 < p ≤ 0.05, **, 0.001 < p ≤ 0.01; ***, 0.0001 < p ≤ 0.001; **** p ≤ 0.0001. Scale bars: 500μm.

### SB increases fat accumulation, at least in part, through the SKN-1/Nrf2 signaling pathway

SKN-1 (an Nrf2 homolog) is a well-known pathway associated with lipid metabolism (Pang et al., 2014; Steinbaugh et al., 2015). SKN-1 activates diverse lipid metabolism genes and reduces fat storage (Pang et al., 2014; Steinbaugh et al., 2015). To assess whether SB treatment increases fat accumulation through SKN-1, *skn-1(zj15)* loss-of-function mutant worms were used. L1-synchronized *skn-1(zj15)* mutant worms were cultured for 72 hours on NGM agar plates with various concentrations of SB and 0.016% glucose. Fat accumulation was measured by ORO staining. While the overall fat accumulation levels in *skn-1(zj15)* mutant worms were significantly higher than those in wild-type worms (data not shown) (Pang et al., 2014; Steinbaugh et al., 2015), no significant changes in fat accumulation due to SB and glucose treatment were observed in the *skn-1(zj15)* mutants as compared to vehicle control (**Figures 3A and 3B**). This result indicates that SKN-1 may be involved in SB-induced fat accumulation in wild-type worms.

**Figure 3.**
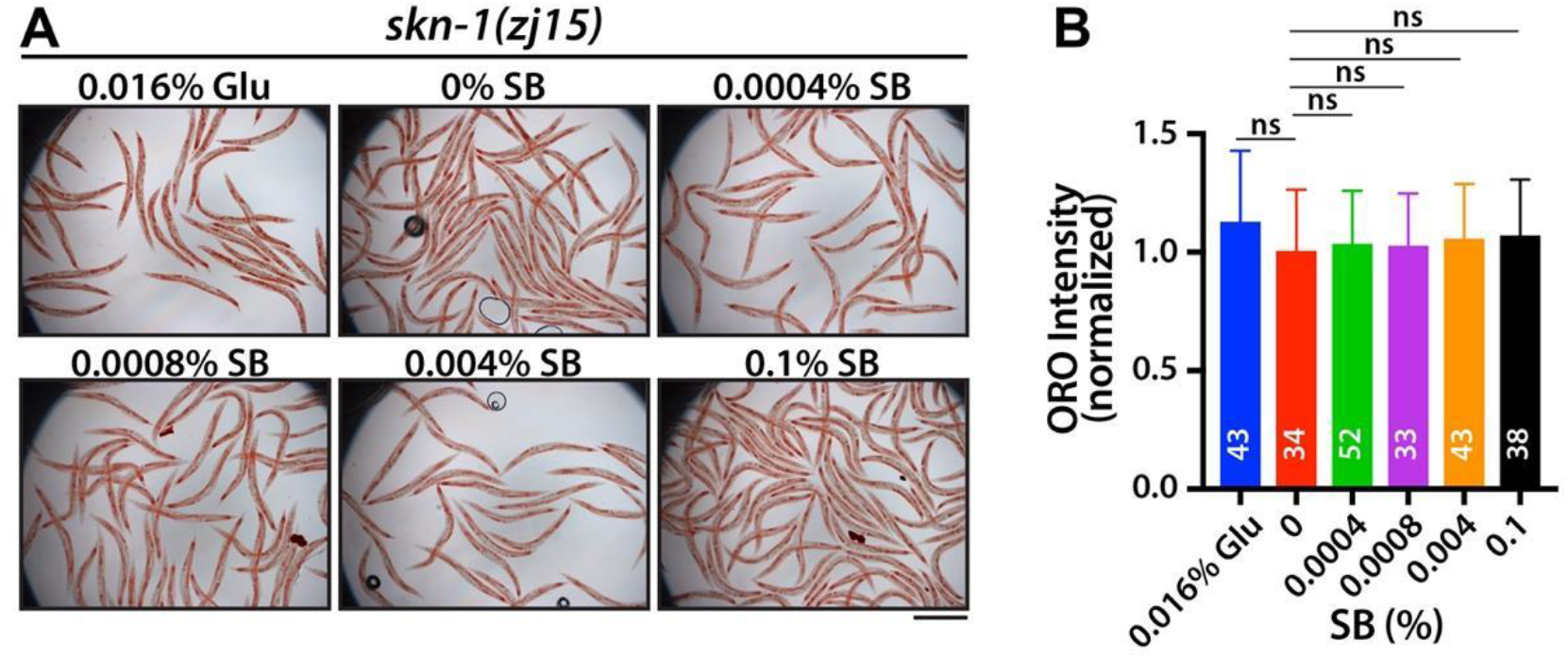
SB-induced fat accumulation may require the SKN-1 signaling pathway. (A) ORO staining with *skn-1(zj15)* mutant worms. (B) Normalized ORO intensity. Not significant (ns), p > 0.05; *, 0.01 < p ≤ 0.05, **, 0.001 < p ≤ 0.01; ***, 0.0001 < p ≤ 0.001; **** p ≤ 0.0001. Scale bar: 500μm.

### SB treatment increases aging primarily through the SKN-1/Nrf2 signaling pathway

The SKN-1 signaling pathway is highly linked to longevity (Ogg et al., 1997; Tullet et al., 2008). Under stress conditions, SKN-1 proteins translocate into the nucleus and activate the expression of their target genes, which are essential for detoxification or stress responses (Ogg et al., 1997; Tullet et al., 2008). To test whether SB treatment affects the longevity of *C. elegans*, L1-synchronized wild-type worms were cultured on OP50-seeded NGM agar plates with various concentrations of SB and 0.016% glucose. Survival rates were recorded daily, and surviving worms were transferred to fresh NGM plates every three days. The average lifespan of wild-type worms was approximately 16.23 ± 3.18 days (**Figures 4A and 4B**). However, treatments with 0.016% glucose and SB significantly reduced the average lifespans of wild-type worms; 15.01 ± 3.69 days for 0.016% glucose, 16.23 ± 3.18 days for 0% SB, 13.27 ± 2.57 days for 0.0004% SB, 15.74 ± 3.75 days for 0.0008% SB, 14.92 ± 3.12 days for 0.004% SB, and 12.98 ± 3.23 days for 0.1% SB (**Figure 4B and Table 1**). Next, to investigate whether SB treatment decreases the lifespan of wild-type worms through the SKN-1 signaling pathway, L1-synchronized *skn-1(zj15)* mutant worms were cultured on OP50-seeded NGM agar plates with various concentrations of SB and 0.016% glucose, and their survival rates were recorded daily as described above. Unlike wild-type worms, *skn-1(zj15)* mutant worms did not show a significant reduction in average lifespan at 0.0004% and 0.0008% SB concentrations. Their average lifespans were only slightly decreased with higher SB treatments (0.004% and 0.1%) (**Figures 4C and 4D**). Specifically, the average lifespans of *skn-1(zj15)* mutant worms were 9.93 ± 2.29 days for 0.016% glucose, 10.21 ± 2.51 days for 0% SB, 10.03 ± 2.35 days for 0.0004% SB, 9.79 ± 2.38 days for 0.0008% SB, 9.23 ± 2.22 days for 0.004% SB, and 9.23 ± 1.67 days for 0.1% SB (**Figure 4D and Table 1**). These results indicate that SB treatment reduces the average lifespan of wild-type worms, at least in part, through the SKN-1 signaling pathway.

**Table 1.** Average lifespans.

| Strain | Treatment | Average lifespan ± (StDv) | p-value (t-test) |
| --- | --- | --- | --- |
| wild-type | 0.016% Glu | 15.01 ± 3.69 days | 0.0420 |
|  | 0% SB | 16.23 ± 3.18 days | - |
|  | 0.0004% SB | 13.27 ± 2.57 days | <0.0001 |
|  | 0.0008% SB | 15.74 ± 3.75 days | 0.3802 |
|  | 0.004% SB | 14.92 ± 3.12 days | 0.0167 |
|  | 0.1% SB | 12.98 ± 3.23 days | <0.0001 |
| *skn-1(zj15)* | 0.016% Glu | 9.93 ± 2.29 days | 0.1667 |
|  | 0% SB | 10.21 ± 2.50 days | - |
|  | 0.0004% SB | 10.03 ± 2.35 days | 0.6873 |
|  | 0.0008% SB | 9.79 ± 2.38 days | 0.3673 |
|  | 0.004% SB | 9.23 ± 2.22 days | 0.0338 |
|  | 0.1% SB | 9.23 ± 1.67 days | 0.0138 |

**Figure 4.**
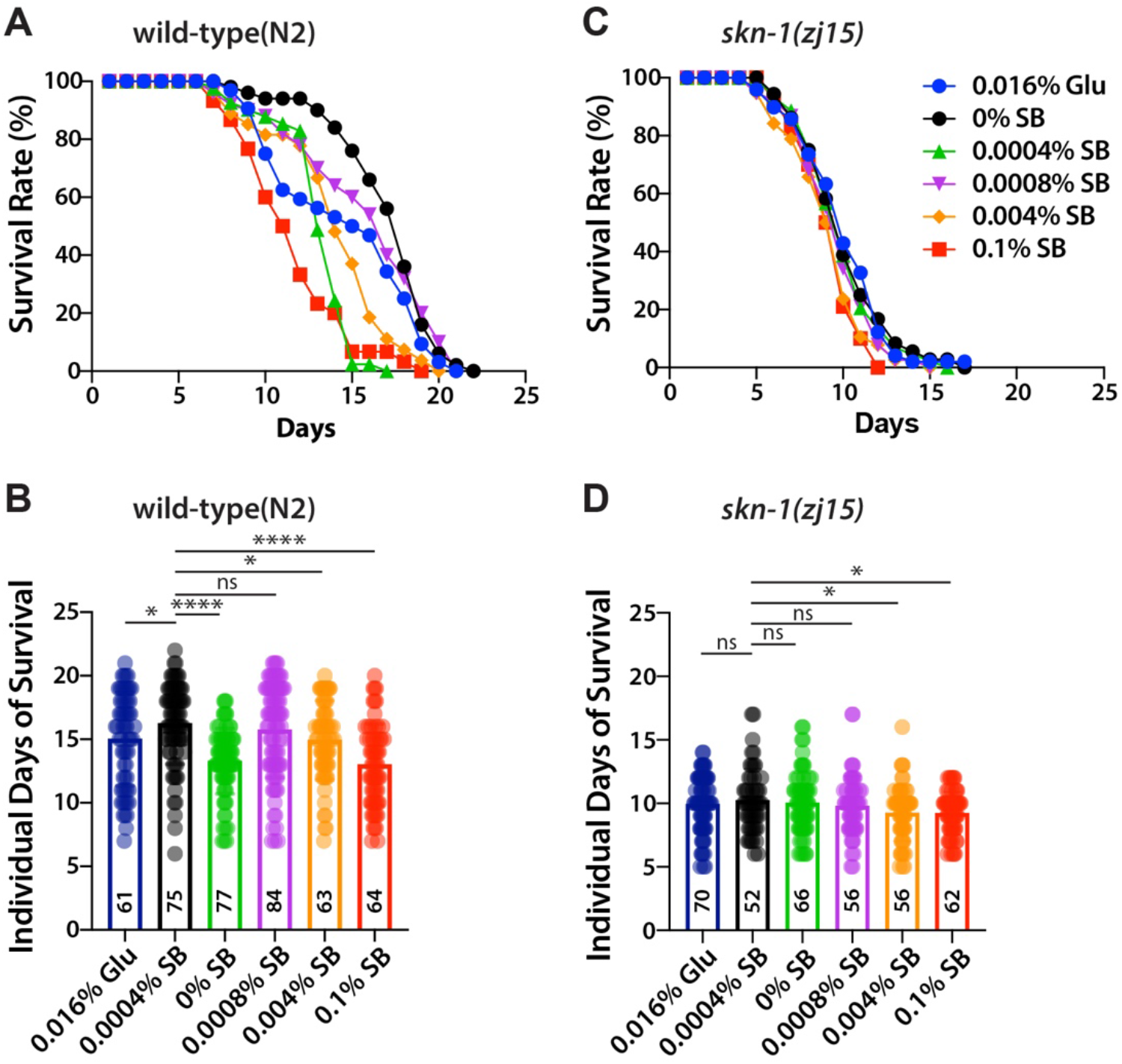
SB shortens the average lifespan through the SKN-1 signaling pathway. (A and C) Lifespan curves of wild-type and *skn-1(zh15)* mutant worms. (B and D) Individual days of survival of wild-type and *skn-1(zh15)* mutant worms. Not significant (ns), p > 0.05; *, 0.01 < p ≤ 0.05, **, 0.001 < p ≤ 0.01; ***, 0.0001 < p ≤ 0.001; **** p ≤ 0.0001. See Table 1 for average lifespans and errors.

### SB inhibits SKN-1 nuclear localization

SB has been known to induce oxidative stress in rodents (Khoshnoud et al., 2018) and zebrafish (Gaur et al., 2018). The major regulator of the oxidative stress response is SKN-1. Under oxidative stress conditions, SKN-1 proteins translocate into the nucleus and increase the expression of detoxification and antioxidant-related genes, thereby reducing cellular oxidative stress (Inoue et al., 2005). To test whether SB could inhibit the nuclear localization of the SKN-1 protein, transgenic worms expressing a *skn-1(b/c)::gfp* gene were employed. SKN-1 proteins are predominantly localized in the cytoplasm under normal conditions (**Figure 5A** left), but they translocate to the nucleus in the intestine (**Figure 5A** right) in response to oxidative stress from various sources, such as sodium arsenite (mitochondrial toxin). The *skn-1(b/c)::gfp* transgenic worms were cultured on OP50-seeded NGM agar plates with various concentrations of SB and 0.016% glucose at 20°C. 48 hours later, the transgenic worms were collected from the plates with M9 buffer and exposed to 5 mM sodium arsenite for 30 minutes. The nuclear localization of SKN-1::GFP proteins in the intestines was analyzed by staining using antibodies against GFP and nuclear pore complex (NPC) proteins. The ratio of SKN-1::GFP proteins in the nucleus to the cytoplasm was measured using ImageJ software. While SKN-1::GFP proteins exhibited dramatic translocation into the nucleus upon treatment with 5 mM sodium arsenite in the absence of SB, however, SKN-1 nuclear localization was significantly inhibited in the presence of SB even at lower concentrations, including 0.0004% (**Figure 5B**).

**Figure 5.**
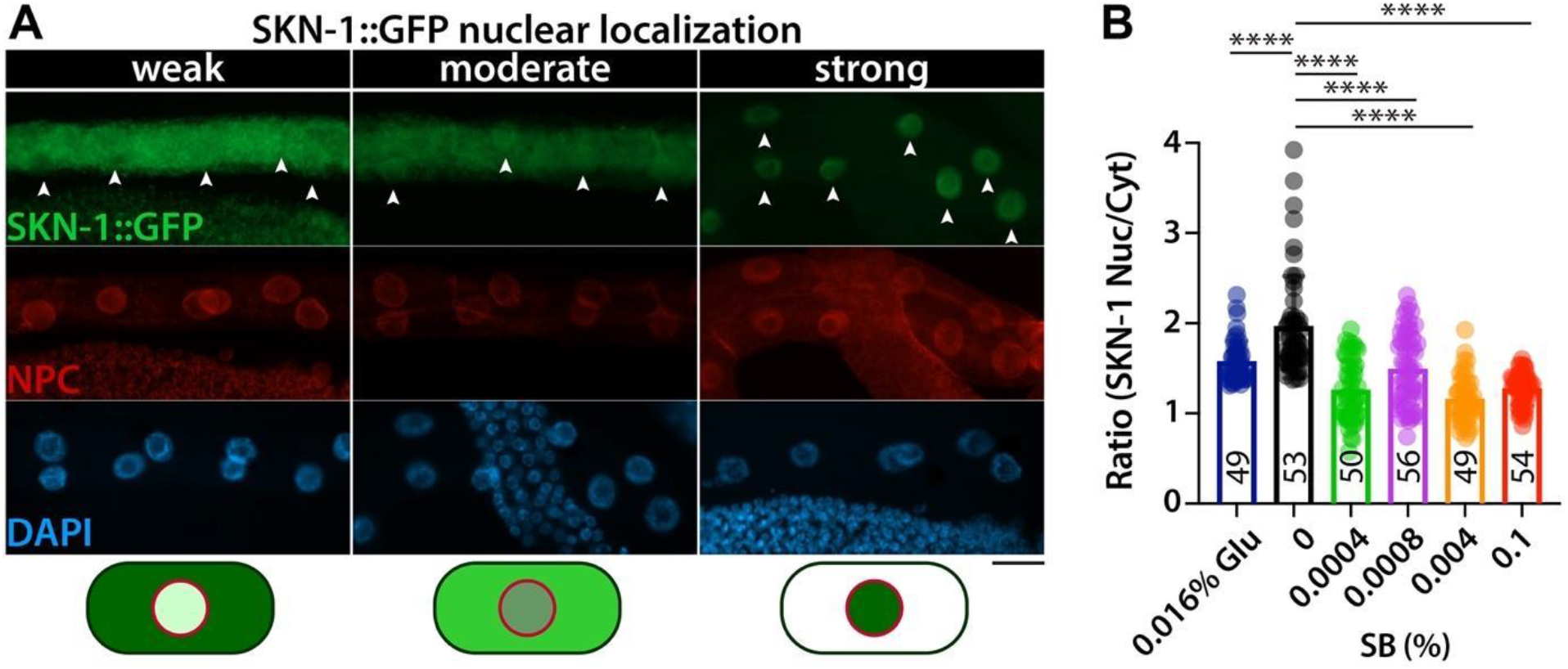
SB inhibits SKN-1 nuclear localization under oxidative stress conditions. (A) Immunostaining. Dissected intestines were stained with antibodies against GFP and the nuclear pore complex (NPC) proteins. All images were acquired using the same setting and magnification. Scale bar: 20μm. (B) The ratio of GFP intensities in the nucleus to the cytoplasm. Below are the images of SKN-1::GFP (green) dynamics and NPC proteins (red). Not significant (ns), p > 0.05; *, 0.01 < p ≤ 0.05, **, 0.001 < p ≤ 0.01; ***, 0.0001 < p ≤ 0.001; **** p ≤ 0.0001.

## DISCUSSION

In this study, we aimed to explore the effect of SB on obesity and aging using the nematode *C. elegans* as a model system. Our results reveal that SB treatment increases fat accumulation and aging, at least in part, through the SKN-1 signaling pathway (**Figure 6**). These findings suggest potential risks associated with SB consumption in the context of obesity and aging.

**Figure 6.**
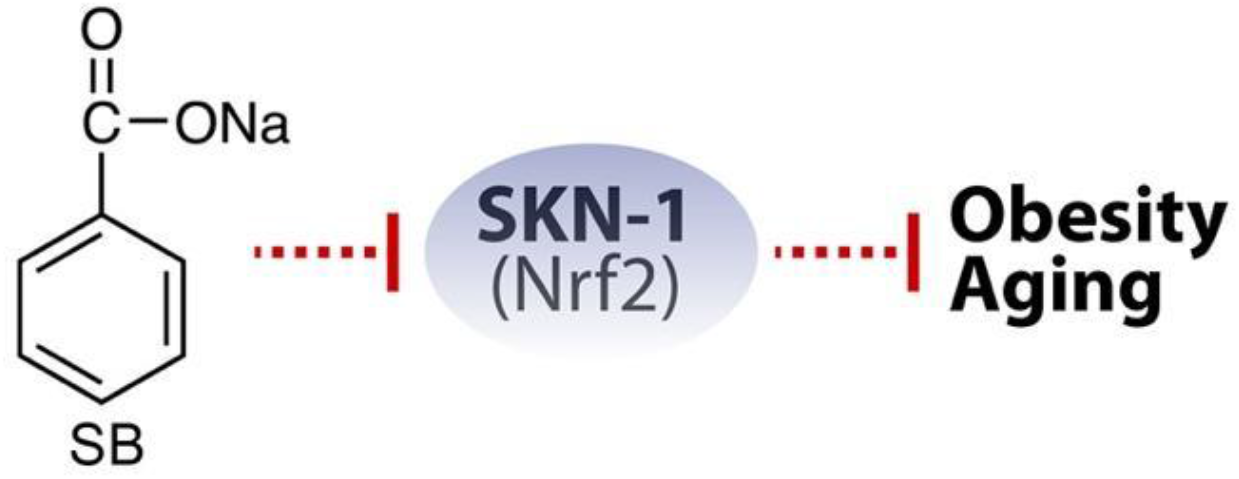
A potential action model of SB in obesity and aging.

SB has been known to inhibit the growth of bacteria, fungi, and yeast. A recent study demonstrated that SB slows the larvae’s developmental timing and shortens the adult lifespan in *Drosophila melanogaster* by affecting endocrine hormone levels and commensal microbiota (Asejeje et al., 2023; Dong et al., 2022). SB has also been associated with reduced reproductive capacity, developmental defects, oxidative stress, and anxiety-like behavior (El-Shennawy et al., 2020; Gaur et al., 2018). In zebrafish, SB treatment induces delayed hatching, morphological defects, oxidative stress, and abnormal behaviors in developing larvae (Gaur et al., 2018). However, in *C. elegans*, SB treatment did not lead to significant developmental defects or reproductive problems, even at a high concentration (0.1%) of SB (data not shown). Questions persist regarding the potential occurrence of these abnormalities in *C. elegans* over multiple generations with long-term SB exposure.

We show that SB treatment increases oxidative stress by disrupting the SKN-1-mediated stress response mechanism (**Figure 5**). Oxidative stress is closely linked to liver diseases such as drug-induced liver injury, viral hepatitis, and alcoholic hepatitis (Zhou et al., 2022). SB exposure affects the lipid profile and parameters of liver and kidney functions in rats (Walczak-Nowicka and Herbet, 2022). Furthermore, SB has been shown to induce damage to kidney structure and function (Zeghib and Boutlelis, 2021). Specifically, SB elevates hepatic oxidative stress and inflammation in liver injury (Nezu and Suzuki, 2020). Nrf2 plays a critical role in various liver diseases, including metabolic dysfunction-associated fatty liver disease (Zhou et al., 2022), and is considered a potential therapeutic target for non-alcoholic steatohepatitis (Galicia-Moreno et al., 2020). Nrf2 also contributes to protecting the kidney from oxidative damage (Nezu and Suzuki, 2020). Thereby, SB may affect the functions of the liver and kidney and their associated diseases, most likely through the modulation of the Nrf2 signaling pathway.

We investigated the effect of SB on fat accumulation and aging in wild-type worms using five different concentrations (0%, 0.0004%, 0.0008%, 0.004%, and 0.1%). The effect of SB on fat accumulation and aging was high at 0.0004% and 0.1% SB. However, at 0.0008% and 0.004%, the effects were lower compared to 0.0004%. It is interesting that SB induced more significant fat accumulation in relatively low dosage, which is consistent with what was observed in mice (Olofinnade et al., 2021). While a 0.0125% SB diet significantly increased body weight and food intake, higher SB concentrations (0.025% and 0.05%) did not alter body weight compared to control groups (Olofinnade et al., 2021). It was proposed that SB may enhance food palatability at a 0.0125% concentration (Olofinnade et al., 2021), but higher concentrations could potentially reduce food intake (Nair, 2001). To explore the potential impact of SB on food intake in *C. elegans*, we evaluated the pharyngeal pumping rate. Our findings revealed a significant increase in pharyngeal pumping rate at higher concentrations (≥ 0.004%) of SB (data not shown), indicating that food intake was not a major factor of the more pronounced fat accumulation and aging effects observed in the 0.0004% dosage group.

In *C. elegans*, lipid metabolism and longevity are also regulated through an insulin signaling pathway, including DAF-2 (an IGF homolog) and DAF-16 (a FOXO homolog) (Hellerer et al., 2007). In *C. elegans*, DAF-2 has been known to inhibit both DAF-16 and SKN-1 through AKT-1/2 kinases in response to external stress (Cahill et al., 2001; Hertweck et al., 2004; Lin et al., 2001). Both DAF-16 and SKN-1 are involved in the oxidative stress response and longevity (Ogg et al., 1997; Tullet et al., 2008). However, DAF-16 and SKN-1 exhibit opposite functions in lipid metabolism – DAF-16 promotes fat accumulation, whereas SKN-1 inhibits it (Pang et al., 2014; Xu et al., 2019). A recent study also demonstrated that SKN-1 functions as a negative regulator of DAF-16 (Deng et al., 2020). Our unpublished results revealed that SB treatment inhibited the expression of the *daf-2* gene. Therefore, we propose that more complicated regulatory networks, including SKN-1 and insulin signaling pathways, may be involved in SB-induced fat accumulation and aging. Although the detailed mechanism by which SB affects the insulin signaling pathway remains unclear, future research addressing this aspect will contribute to a better understanding of SB-induced obesity and aging, thereby significantly advancing our knowledge in this field.

## ACKNOWLEDGEMENTS

We are grateful to Dr. Lee at ECU for *C. elegans* strains and antibodies against GFP and NPC and for helpful discussions during our work. QV225 [*skn-1(zj15)*] and LD1 [*idIs7 skn-1b/c::GFP + rol-6(su1006)*] strains were obtained from the Caenorhabditis Genetic Center (CGC), which is supported by the NIH-National Center for Research Resources.

